# A new skin supplement Purasomes XCELL: an integrative approach in regenerative aesthetics

**DOI:** 10.1101/2025.01.08.631888

**Authors:** Antonio Salvaggio, Federica Coppa, Maria Violetta Brundo

**Affiliations:** Department of Biological, Geological and Environmental Science, University of Catania, Catania

**Keywords:** Bovine colostrum, *Astragalus membranaceus*, exosomes, telomeres, Telomere Analysis Technology, fibroblasts

## Abstract

Food supplements are taking on an increasingly important role as potential support tools for well-being and health. Supplements that offer essential nutrients and bioactive compounds can help fill nutritional gaps and provide specific benefits to complement health and wellness promotion strategies. Many supplements contain bioactive compounds used to enhance skin health and appearance, but only few biomolecules influence telomere lengths. Aims of this study was evaluated *in vitro* effectiveness of a new supplement Purasomes XCELL that contains natural compounds extracted from bovine colostrum and *Astragalus*, on skin aging using telomere length as an aging biomarker. Our results suggest that Purasomes XCELL has protective effect that is manifested already 4 weeks of treatment, but the telomere shortening rates on human fibroblasts were even more pronounced at 8 weeks. This new supplement could be highly effective on skin aging.

## 1. Introduction

Aging is an inevitable and individual physiological process and the way in which we age varies depending on the social, economic and cultural context in which we live, but also on the genetic characteristics of the individual [1]. In the last decade, numerous studies have focused on the search for molecular biomarkers and on the mechanisms involved in biological aging, in order to slow down this process and maintain a good quality of life [2, 3]. Among the various proposed biomarkers are telomeres, stretches of highly repeated DNA located at the ends of each chromosome arm of eukaryotic DNA [4-6]. Telomeres preserve the structural integrity of DNA during each replication cycle, as DNA polymerase is unable to replicate the chromosome until it is terminated. Numerous studies have hypothesized that the progressive shortening of telomeres with each replicative cycle is associated with cellular aging (senescence phase). They, therefore, act as a molecular timer controlling the lifespan of a eukaryotic cell [4-6]. The structure of telomeres is made up of non-coding nitrogenous bases (5’-TTAGGG-3’). However, recent studies have shown that the telomeric structure produces transcripts called TERRA, which are hypothesized to be involved in the regulation of telomerase [7,8]. In mammals, telomeres are highly conserved, and in humans the length of the telomere segment is between 5,000 and 15,000 base pairs. This long stretch of repetitive DNA sequences is characterized by a single-stranded protrusion at the 3’ end, which threads into the end of the chromosome, creating a conformation called a T-loop, stabilized by a group of proteins (Shelterin complex). Telomeres are closely linked to cellular aging, especially in dermal cells. Telomere shortening in skin fibroblasts may lead to epidermal aging and barrier function defects [9, 10]. Telomere length is regulated by the enzyme telomerase through the addition of guanine-rich repetitive sequences [11]. Telomerase is active only in germ cells and stem cells, but not in somatic cells [12]. In recent decades, several studies have shown that biomolecules (such as immunoglobulins, growth factors and cytokines) and exosomes present in bovine colostrum are involved in various cellular metabolic mechanisms and, having antioxidant properties, can help prevent cellular damage and slow down the signs of aging [13-15]; also astragalus, a plant widely used in traditional Chinese medicine, has garnered significant attention for its potential to activate telomerase and extend telomere length, making it a promising natural nutritional supplement anti-aging [16].

Objective of this study is to demonstrate the protective effect of a new supplement that contains active biomolecules purified from bovine colostrum and *Astragalus membranaceus* extract on skin aging using telomere length as an aging biomarker.

## 2. Materials and Methods

For our research we purchased a supplement Purasomes XCELL that utilize a new technology called AMPLEX plus, containing exosomes purified from bovine colostrum passively loaded with growth factors and cytokines purified from bovine colostrum. Purasomes XCELL contains also *Astragalus membranaceus* extract, blue Icelandic spirulina, organic Moldavian dragonhead and blue spirulina derived vitamin B12.

### 2.1 Propagation and maintenance of cells

Normal adult human dermal fibroblasts (HDFa) (ThermoFisher Scientific) were cultured under standard conditions and were used with negative control. In particular, cells were cultured in Dulbecco’s Modified Eagle Medium (DMEM) supplemented with 10% FBS or 5% FBS, streptomycin (0.3 mg mL^−1^) and penicillin (50 IU mL^−1^), and GlutaMAX (1 mM) using 75 cm^2^ polystyrene culture flasks. Cells were maintained in a humidified environment (37 °C and 5% CO_2_) and split every 2-3 days depending on cell confluence. Three different concentrations of AMPLEX plus (0.125%, 0.25%, 0.5%) were added to the primary culture medium (5% and 10% FBS) to test for telomere shortening rate. As a positive control, cells were treated with 80 μM of H_2_O_2_. Fresh treatments were added to the primary culture at each passage when cells were split. In total, were analyzed nine groups of cells for eight weeks of treatment.

Cell growth was monitored for each treatment group by counting cells at each passage. Proliferative capacity (PC) of cells was calculated with the formula PC = (log (Nn/Nn-1))/log 2 where n is the passage number and N is the number of cells. One PC was equivalent to one round of cell replication [17].

### 2.2 Telomere shortening rate assay

Telomere length was measured by a Telomere Analysis Technology (TAT) according to [18]. Three distinct variables were determined during TAT analysis: the median telomere length, 20^th^ percentile telomere length, and the percentage of telomeres < 3 kilobase pairs (kbp). All telomere length measurements were expressed as the number of base pairs (bp).

### 2.2 Statistical analysis

Data were reported as the mean of at least three independent experiments. Telomere length distribution and median telomere length were calculated by inhouse software.

## 3. Results and Discussion

Colostrum, the first milk produced by humans and other mammals in the earliest days after giving birth, is rich of nutrients, but also of bioactive components with immune enhancing properties, such as immunoglobulins, lactoferrin, lysozyme, lactoperoxidase, α-lactalbumin, β-lactoglobulin [19]. The concentration of these compounds is variable and depends on many factors, including breed, productivity, parity, feeding intensity, season of the year, and/or production system [20, 21]. Colostrum contains high concentrations of growth factors, that have an important role in fundamental cellular processes, including growth, proliferation, differentiation, and survival. They binding to specific receptors on the cell surface, initiating a cascade of intracellular signaling events that stimulates the regulation of gene expression and cellular functions [22]. The two main growth factors, insulin-like growth factors 1 and 2 (IGF-1 and IGF-2) and transforming growth factors alpha and beta (TGF-α and TGF-β), are only found in colostrum [22]. Bovine colostrum is a unique and powerful source of these development factors since it naturally contains them, unlike other supplements or synthetic substitutes [23, 24]. In Purasomes XCELL, the bioactive molecules of colostrum are purified, concentrated and are partly conveyed through exosomes which are passively loaded with these biomolecules (AMPLEX plus technology). In recent years, exosomes have attracted the scientific community as nanovesicles of endosomal origin, which can be secreted by a variety of cells and have the same characteristics as progenitor cells. To date, the administration of exosomes orally is the subject of attention because it represents a non-invasive and efficient method for delivering bioactive molecules into the intestine. Different reports show that exosomes can resist the degradative conditions of the gastrointestinal tract after oral administration, accumulating regionally in the intestine, where they are absorbed for systemic biodistribution [25].

Another component present in EXCEL, is *Astragalus membranaceus*. At present, more than 200 compounds have been isolated and identified from Astragalus, such as many biological activities molecules [26, 27]. *Astragalus* has multiple pharmacological effects, such as anti-oxidative-stress, anti-inflammatory, immuno-regulation, vascular protective effects, anti-neurodegeneration, and anti-aging effects being able to activate numerous via of cellular signaling pathways. Specifically, its applications in aging and aging-related diseases, were showed in many studies [16, 26-28].

In order to demonstrate the protective effect of Purasomes XCELL on skin aging we used *in vitro* tests. Proliferative capacity of fibroblasts progressively increased in cells treated with XCELL compared to untreated cells, over the eight weeks. (Figure 1). The increase is particularly evident in cultures incubated with 5% FBS + 0.5% of XCELL, compared to those incubated with 10% of FBS, due to osmolarity problems. This suggests that the addition of the components present in the supplement increases proliferation rates of human fibroblasts.

**Figure 1.**
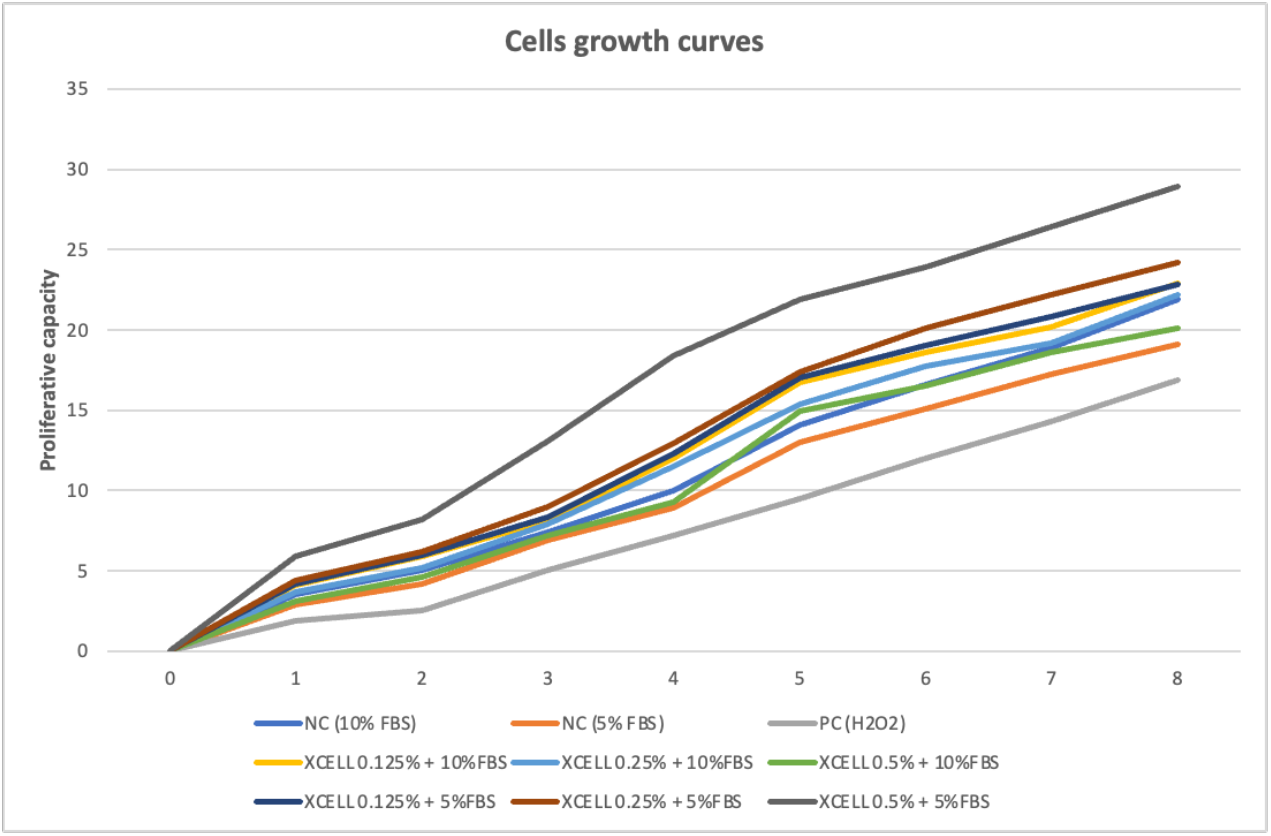
Proliferative capacity of fibroblasts after eight weeks of treatment. NC, negative control (10% and 5% of FBS); PC (80 μM of H_2_O_2_). Cells were treatment with 0,125%, 0,25% and 0,5% of XCELL supplement in10% and 5% of FBS. Fresh treatments were added to the primary culture at each passage when cells were split, every 2-3 days.

As shown in Figure 2, telomere length continued to decrease in cells throughout the study period. Median telomere lengths in the negative control progressively decreased from week 4 to week 8, but in XCELL-treated cells, these changes are less evident, suggesting that the treatment with XCELL not are associated with a notable decrease in telomere shortening after 4/8 weeks of culture. In fibroblasts treated with 0,5% of XCELL + 5% of FBS, in fact telomere length was comparable to value of start experiment (0 week).

**Figure 2.**
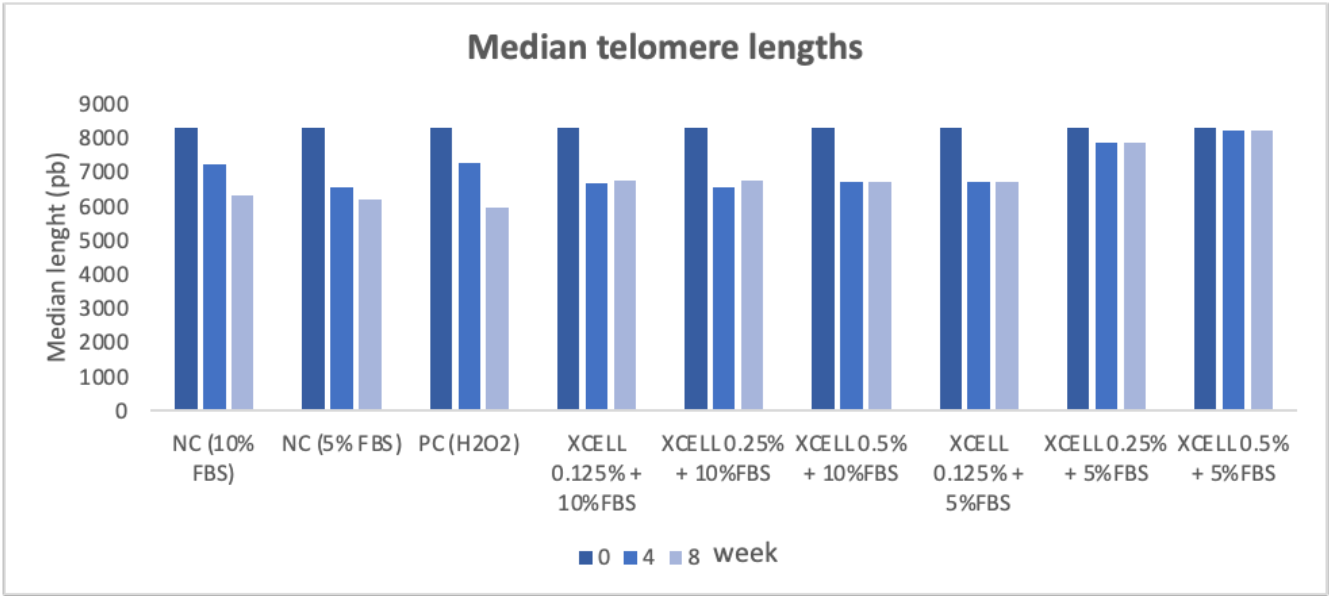
Median telomere lengths obtained with Telomere Analysis Technology. NC, negative control (10% and 5% of FBS); PC (80 μM of H_2_O_2_). Fibroblasts were treatment with 0,125%, 0,25% and 0,5% of XCELL supplement in10% and 5% of FBS. Results at 0, 4 and 8 weeks.

Figure 3 shows the 20^th^ percentile telomere lengths. Negative control samples decreased progressively over the 8 weeks, instead, in the XCELL-treated groups (Figure 3), telomeres are longer, thus demonstrating a protective effect. The 20^th^ percentile indicates in fact the telomere length below which 20% of the observed telomeres fall. As such it is an estimator of the percentage of short telomeres in the cells.

**Figure 3.**
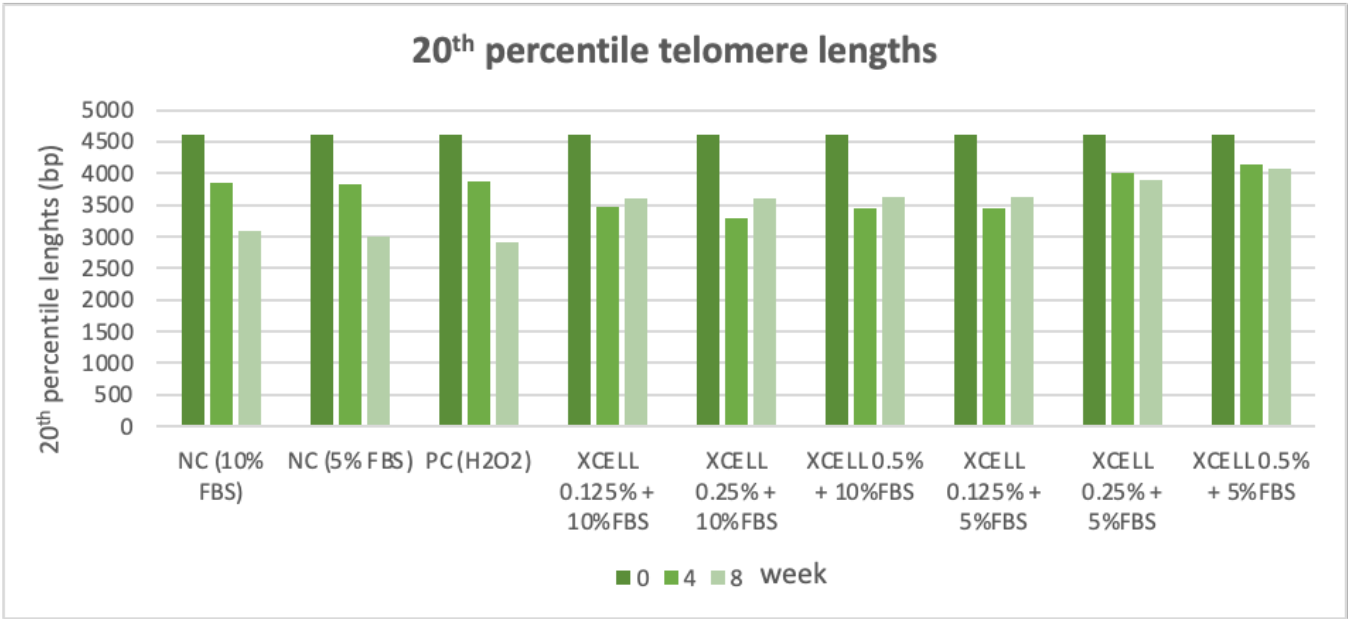
20^th^ percentile telomere lengths obtained with Telomere Analysis Technology. NC, negative control (10% and 5% of FBS); PC (80 μM of H_2_O_2_). Fibroblasts were treatment with 0,125%, 0,25% and 0,5% of XCELL supplement in10% and 5% of FBS. Results at 0, 4 and 8 weeks.

Figure 4 shows the percentage of telomeres less than 3 kbp. For both controls, percentages increased during the 8 weeks. In treated cells instead there was only a small increase in percent telomere <3kbp during of the 8 weeks, suggesting that the presence of XCELL appears to exert a protective effect.

**Figure 4.**
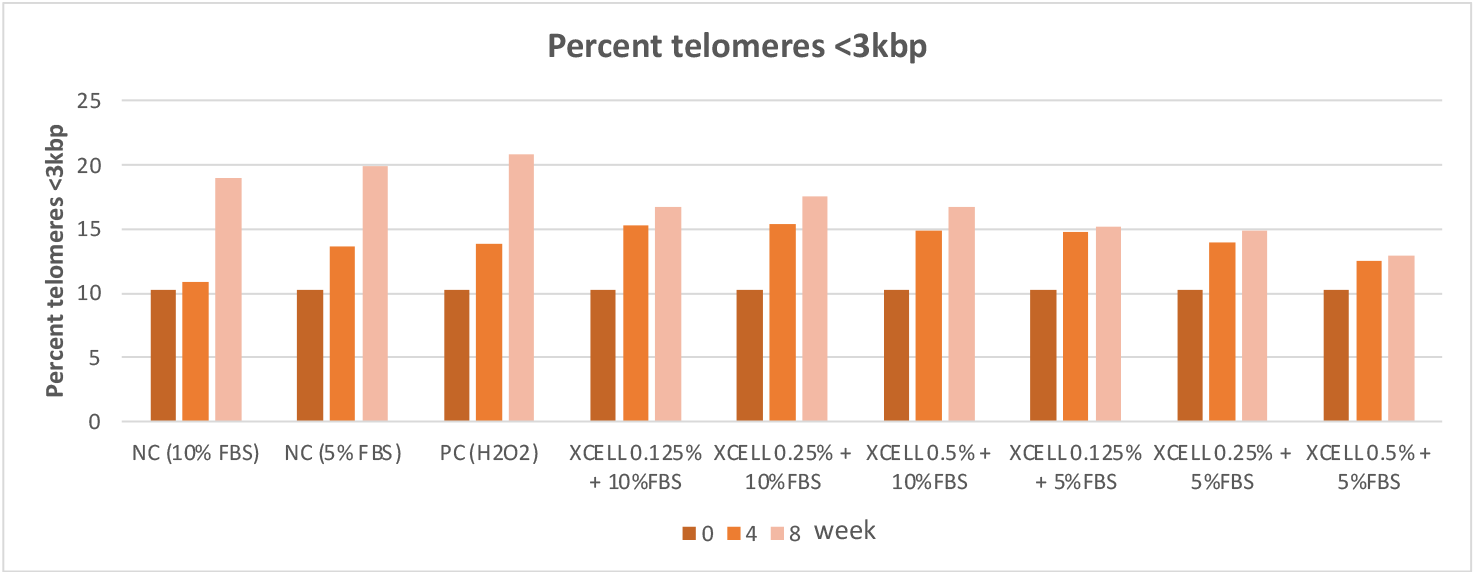
Percent telomere <3kbp obtained with Telomere Analysis Technology. NC, negative control (10% and 5% of FBS); PC (80 μM of H_2_O_2_). Fibroblasts were treatment with 0,125%, 0,25% and 0,5% of XCELL supplement in10% and 5% of FBS. Results at 0, 4 and 8 weeks.

Our results suggest that XCELL was protective effect is manifested already 4 weeks of treatment, but the telomere shortening rates on human fibroblasts were even more pronounced at 8 weeks. One of the active ingredients of XCELL is a mixture of bioactive factors (immunoglobulins, growth factors, cytokines) purified from bovine colostrum and passively loaded into exosomes extracted from bovine colostrum (AMPLEX plus technology). Many studies suggests that bovine colostrum contains a numerous component that play various roles, also anti-aging benefits [14, 24, 29, 30]. As we understand more about colostrum, we recognize its far-reaching potential in increasing longevity, and daily supplementation with biologically active bovine colostrum offers certainly benefits many people [31]. Also, the other active ingredient in XCELL, *Astragalus membranaceus* extract, due to its telomerase-activating compounds, has shown benefits *in vitro* and *in vivo* experiments [32]. Recently, also some human trials have produced satisfactory results [33], in fact the Astragalus contains TA-65, a small molecule telomerase activator that was discovered in a systematic screening of natural product extracts from traditional Chinese medicines [34, 35].

## 4. Conclusion

Many cosmeceuticals contain bioactive compounds used to enhance skin health and appearance. The difference is that XCELL is a natural source of these compounds and can be considered a nutraceutical, namely nutritive food possessing good functional and medicinal value, whose intake prevents aging and lifestyle disorders. The nutritional supplements derived from natural resources, such as XCELL are presently of increasing demand to maintain the structure and function of the body [24, 33, 36, 37].

## Author Contributions

Conceptualization, A.S. and M.V.B.; methodology, F.C.; software, F.C.; validation, F.C.; formal analysis, F.C.; data curation, A.S.; writing-original draft preparation, A.S. and M.V.B.; writing-review and editing, A.S., and M.V.B.; supervision, A.S. and M.V.B.; funding acquisition, M.V.B. All authors have read and agreed to the published version of the manuscript.

## Institutional Review Board Statement

This study was performed in line with the principles of the Declaration of Helsinki and does not require approval by the Ethics Committee of University of Catania.

## Informed Consent Statement

Not applicable.

## Data Availability Statement

The data presented in this study are available on request from the corresponding author.

## Acknowledgments

S.P. thanks the Ph.D. program.

